# HISTOLOGICAL EVALUATION OF THE EFFICACY OF COLLAGEN MEMBRANE IMPREGNATED WITH RECOMBINANT HUMAN FIBROBLAST GROWTH FACTOR-2 (rhFGF-2) FOR GINGIVAL RECESSION DEFECTS IN BEAGLE DOGS

**DOI:** 10.1101/2025.10.30.685538

**Authors:** R Viswa Chandra, Kidambi Sneha, S Anoop Puthoochirayil, V Sthevaan

## Abstract

**BACKGROUND:** One of the most common esthetic concerns associated with the periodontal tissues is gingival recession. Many techniques have been introduced to treat gingival recession, including connective tissue grafting, or CTG; various flap designs; GTR membranes. However, their combination with biologic modifiers has not been explored. So, The present animal study aims to histologically compare collagen membrane impregnated with recombinant human fibroblast growth factor-2 (rhFGF-2) with that of plain collagen membrane in the treatment of Miller’s Class I and II gingival recessions for root coverage and healing process in beagle dogs.

**MATERIALS AND METHODS:** 18 beagle dogs of ages between 4-6 years were included in the study. The study was conducted as split mouth study on test site collagen membrane impregnated with recombinant human fibroblast growth factor-2 (rhFGF-2) was placed, on control site plain collagen membrane was placed. Histologically bone regeneration, mineralized tissue volume, membrane remnants and clinically recession depth were analysed at baseline, 3 months and 6 months.

**RESULTS:** No significant difference was observed between the test and the control groups from baseline to 3 months *(p=0*.*331 NS)* bone regeneration, whereas, significant difference was observed from baseline to 6 months *(p=0*.*008*)*. The three months histological specimen of the test and control sites showed woven bone (NB) with rich vasculature along with abundant collagen bundles. Remnants of the membrane can also be observed at the periphery with no signs of multinucleated giant cells. The six months histological report of the test site shows matured bone formation, with rich vasculature, but new bone formation was less in control group. Significant difference was seen between test and control groups for mineralized tissue formation at 6-months. *(p=0*.*003)*. Highly significant difference was seen for membrane degradation in test group *(p≤0*.*001**)* compared to control group. Significant differences were observed between the test and the control groups for the root coverage from baseline to 3 months. *(p=0*.*008*)* and from baseline to 6-months highly significant was observed*(p=0*.*001**)*.

**CONCLUSION:** The three months and six months specimens of the test (rhFGF2 membrane) group has shown good amount of bone regeneration and soft tissue healing with limited membrane remnants and reduced recession depth at the test site, when compared to the control group. These results prove that the rhFGF2 growth factor impregnated collagen membrane is highly efficient in treating gingival recession, thus improving the periodontal health.

## INTRODUCTION

Gingival recession is the exposure of root surfaces due to apical migration of the gingival tissue margins; gingival margin migrates apical to the cementoenamel junction.^1^ The level of recession is measured as the distance between the CEJ and gingival margin. Root exposure results in long clinical crowns which is not esthetically pleasing and may eventually lead to root caries and sensitivity.^2^

Growth factors have an essential role in healing process and tissue formation, repair, chemotaxis and cell proliferation. Several bioactive molecules such as PDGF, TGF-1, BMP-2 EMD have shown positive results in stimulating periodontal regeneration.^3^ Resorbable collagen barriers have been used clinically for guided tissue regeneration procedures; however, their combination with biologic modifiers has not been explored.

Fibroblast growth factor (FGF2) has ability to induce proliferation in undifferentiated cells.^4^ FGF-2 also showed potent angiogenic ability and mitogenic activity on mesenchymal cells. Recently, in intra-bony defects rhFGF-2 has showed regeneration of periodontal tissues.^5^ rhFGF-2 stimulates mesenchymal cells to differentiate into cementoblasts, osteoblasts, and collagen-forming cells.^6^ Collagen-based biomaterials are commonly used as delivery vehicles for protein drugs, including FGF-2, because they can form a stable polyanionic complex with FGF-2. ^7^

In a preliminary study, a cross-linked collagen membrane was utilized as a carrier for rhFGF-2 for Miller’s class I and II recession defects. Results of this study concluded that rhFGF-2-impregnated collagen membranes showed significant results in terms of increasing the width of keratinised gingiva and recession coverage.^8^ However, subsequent trials with more focus of clinical efficacy with regard to healing through histological analysis maybe more beneficial.

Among the animal models, for periodontal study dogs are one of the commonly chosen models because of the high occurrence of periodontal disease and the similar etiologic factors as humans. Periodontal diseases in the dogs closely mimic the disease in humans. As with humans, gingival recession is an outstanding feature in dogs with periodontitis.^9^ Dogs also served as the animal model in muco-gingival surgery by creating recessions on the canines to study wound healing. Hence, the most commonly used animal model in periodontal research seems to be beagle dogs due to reproducible critical-sized defects. ^10^

The present animal study aims to histologically compare collagen membrane impregnated with recombinant human fibroblast growth factor-2 (rhFGF-2) with that of plain collagen membrane in the treatment of Miller’s Class I and II gingival recessions for coverage and healing process in beagle dogs.

## MATERIALS AND METHODS

### Participants and Sample

The animal ethical committee clearance was obtained for this study with the study code of SVS/AEC/204. This study followed the guidelines of CPCSEA(Committee for the Purpose of Control and Supervision of Experiments on Animals)

18 beagle dogs (*Canis lupus*) of ages between 4-6 years were included in the study. All dogs were fed a dry coarse diet, which is uniformly changed for the whole group. It was conducted as split mouth study on test site collagen membrane impregnated with recombinant human fibroblast growth factor-2 (rhFGF-2) was placed, on control site plain collagen membrane was placed.

### Outcomes Measured

#### New Bone Formation

Newly formed bone was assessed based on the histologic scoring method for bone regeneration by Han et al, where the scores are as follows: 0, no evidence of newly formed bone; 1, ≤ 10% of the original bone defect; 2, ≤ 20% of the original bone defect; 3, ≤ 30% of the original bone defect; 4, ≤ 40% of the original bone defect; 5, ≤ 50% of the original bone defect; 6, ≤ 60% of the original bone defect; 7, ≤ 70% of the original bone defect; 8, ≤ 80% of the original bone defect; 9, ≤ 90% of the original bone defect; and 10, ≤ 100% of the original bone defect. *(Figure 4)*

#### Mineralized Tissue Volume Analysis

From operative sites of each section, ten regions of interest (ROIs) per slide were imaged (*Olympus BX53® microscope, DSS Group, New Delhi, India*) at 40X magnification. The mineralized tissue volume (MTV) was assessed as follows; The images were opened in ImageJ® and are converted into black and white. The “wand tool” and shift key were used to select the black areas which correspond to the areas of mineralized tissue/ bone. Selecting Analyze>Measure quantifies the mineralized tissue volume (MTV) of the slide which is derived by the formula (mineralized tissue area/total area) *100. The MTV of a group was defined as the average of MTV values for 10 ROIs per slide for each section within a group. *(Figure 5)*

#### Analysing Membrane Remnants

Quantification of membrane degradation was done by application of a classifier that had been generated by a pathologist using the Weka plugin in FIJI. Evaluation of membrane degradation by Weka analysis was performed as follows. In each slide, images were split into tiles of 10× magnification, and the classifier looked for a rounded connective tissue capsule with a clear split from the surrounding tissues suggestive of membrane degradation products.5 The areas of interest were automatically segmented into images. The area occupied by the membrane degradation product was calculated and expressed as (degradation product/total area) × 100 as per a previously reported protocol.*(Figure 6)*

#### Analysing recession depth

Customized resin stents were prepared using room temperature curing resin before the root coverage surgeries for each animal/tooth. The stents were to record recession depths which was the most mid-facial gingival recession from the mesio-distal line at the most coronal gingival level (the flat base of the stents) to the most apical gingival margin. *(Figure 2)*

#### Surgical Procedure

The animals will be anesthetized and will be placed in dorsal recumbency. The operative field will be prepared in a standard manner and the surgical procedures carried out using aseptic methods. Surgical procedure will be carried out as follows: Dehiscence-type gingival recession defects were surgically created bilaterally in canines. Two vertical incisions separated by a distance of 5 mm were made from the gingival margin and extending 7 mm apically in order not to reach the mucogingival line to preserve the limited amount of keratinized gingiva. These incisions were connected apically by a horizontal incision and coronally by an intrasulcular incision. The gingival tissue limited by the incisions was removed using a periosteal elevator.

The exposed intact bone 1 to 1.5 mm coronally from the horizontal incision line was removed by means of bone chisels to expose the root surface 5 mm from the cement-enamel junction (CEJ), and the root surfaces were carefully scaled using hand curettes to remove the root cementum (Hu-Friedy Co., Chicago, IL, USA. Full-thickness flaps were then raised, and the root surfaces were scaled to completely remove the biofilm and the residual inflamed granulation tissue. Reference notches were made using a #1 round bur on the root surface at the base of the defects, at the level of the CEJ and on the crown surface to indicate the precise centre plane of the dehiscence defects and to aid in optimal histologic processing. Bilateral defects were randomly assigned to receive FGF+Collagen or collagen alone. *(Figure 1)*

**Figure 1.**
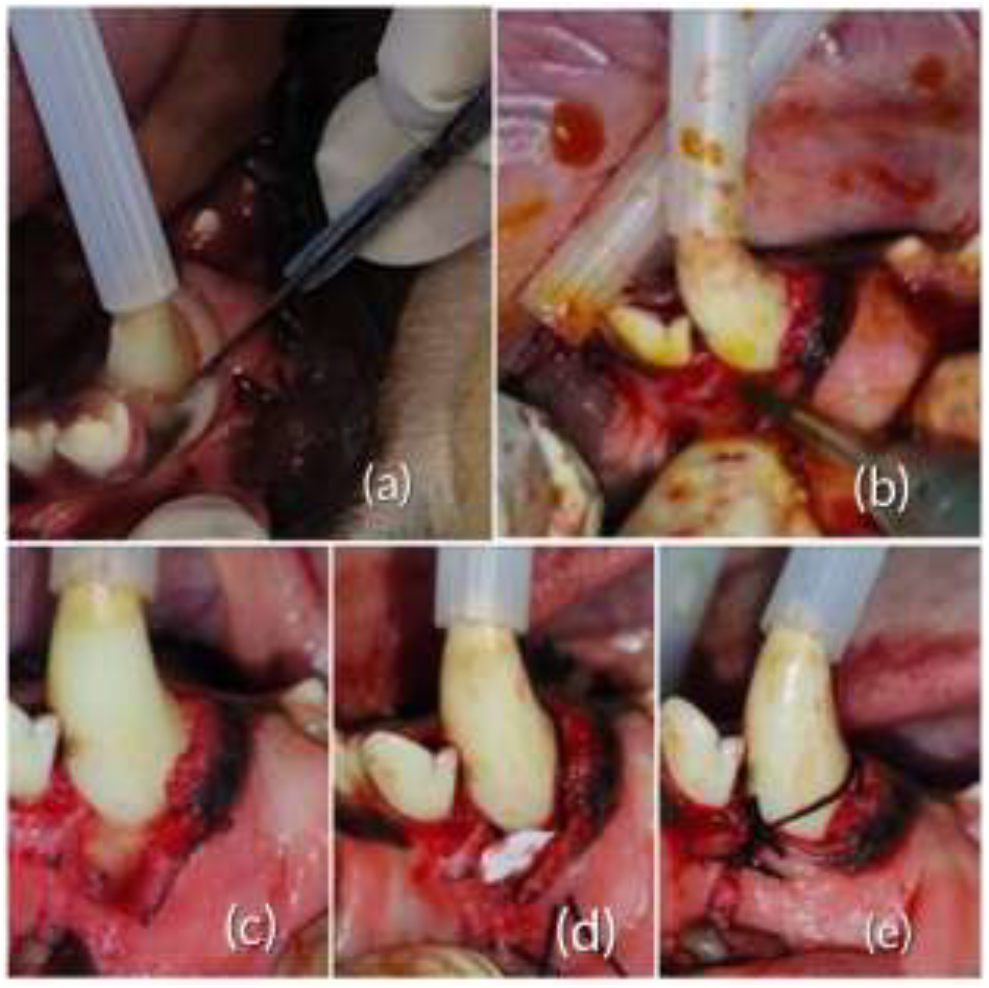
(a) Incision given with no.23 blade to raise the flap; (b) Recession being created with micromotor bur; (c) Recession defect made; (d) FGF 2 membrane placed in the defect; (e) Defect closed with silk sutures

**Figure 2.**
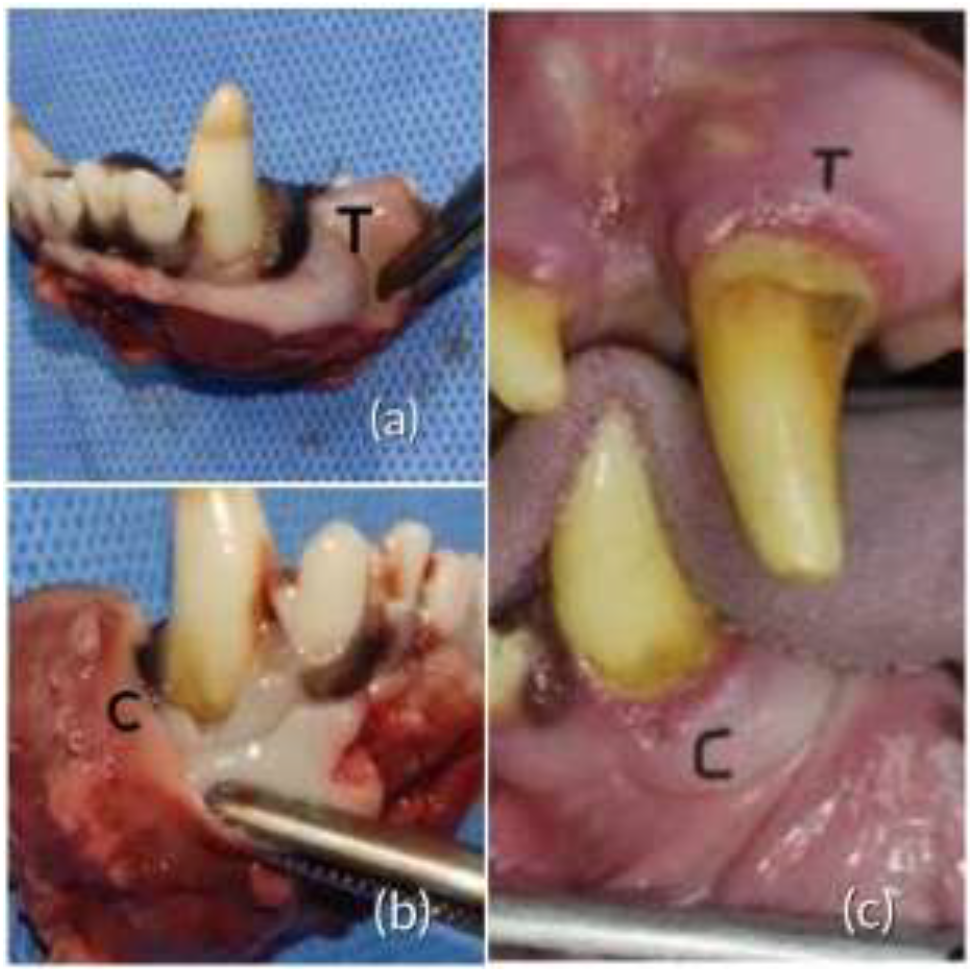
Post operative (a) 3 months post op excised specimen of the test site; (b) 3 months post op excised specimen of the control site; (c) 6 months post operative image of upper-test site, lower-control site

Soon after the surgery, the animals will be caged individually at a temperature of 24 to 28°C and at 70% relative humidity and were kept on a 12 h light/12 h dark cycle. Standard regimen of care will follow. Clinically, at 1month, 3 months and 6 months, the recession coverage is measured from the reference notch at the level of cementoenamel junction and the gingival margin buccally.

#### Euthanasia and Histological Analysis

6 animals each will be sacrificed by a lethal dose of isoflurane at a period of 1, 3 and 6 months respectively. The animals were euthanized by an overdose injection of sodium thiopental. All the defects were dissected along with the surrounding soft and hard tissues. The tissue blocks were fixed in 10% buffered formalin and trimmed. The samples were dehydrated and embedded in polyester resin. The resin blocks were cut bucco-lingually to a thickness of 100 to 150 µm with a low-speed diamond saw. Slides were ground and polished to a final thickness of 35 to 45 µm using a micro grinding system with non-adhesive abrasive discs and stained with toluidine blue.

#### Statistical Analysis

Proportional power calculation was used to determine the sample size and according to the analysis, a minimum 18 subjects/ group when the power of the test is 0.80 at a significance level of 0.05. b data collection: Along with the clinical measurement of height obtained, in histologic specimens, the epithelium healing index by Landry, Turnbull and Howley and the histological scoring method for bone regeneration by Han et al will be the primary tools for measuring histologic healing. Data will be analyzed by using Prism8® GraphPad Software, La Jolla, USA. Measurements will be summarized as median± IQR or mean Â± SD depending on the parameters. Intragroup comparison will be done by Kruskal-Wallis test and intergroup comparison between sites receiving collagen membrane alone and collagen membrane with rhFGF2 will be done by Mann Whitney test or unpaired T test for score data.

## RESULTS

### Intergroup comparison of Bone regeneration at different time-intervals

When the results of bone regeneration were compared within the test group, an increase in bone regeneration was observed from 3 months(5.117±0.70mm) to 6 months (9.312±2.75) with rhFGF2 growth factor impregnated collagen membrane, whereas in control group also there was an increase in bone regeneration from 3 months (5.162±1.105) to 6 months(7.123±1.324), but when the results of bone regeneration were compared between test and control group, no significant difference was observed between the test and the control groups from baseline to 3 months *(p=0*.*331 NS)*, whereas, significant difference was observed from baseline to 6 months. *(p=0*.*008*) (Table 1)(Figure 4)*

**Table 1.** Intergroup comparison of Bone regeneration & Membrane Degradation at different time-intervals.

| Variable | Interval | Test | Control | t-value | p-value |
| --- | --- | --- | --- | --- | --- |
| **Bone regeneration** | *3-months* | 5.117±0.70 | 5.162±1.105 | 0.156 | 0.331(NS) |
|  | *6-months* | 9.312±2.75 | 7.123±1.324 | -0.653 | 0.008* |
| **Membrane Degradation** | *3-months* | 60.19±23.70 | 66.69±14.20 | 12.67 | 0.001** |
|  | *6-months* | 97.54±16.67 | 95.25±16.56 | 9.83 | 0.006* |

*Independent t-test, $p \leq 0.05$ \*(significant), $p \leq 0.001$ \*\* (highly significant), $p \geq 0.05$ NS (Not significant).*

The three months histological specimen of the surgical site (test) shows woven bone (NB) with rich vasculature along with abundant collagen bundles. Remnants of the membrane can also be observed at the periphery with no signs of multinucleated giant cells. The six months histological report of the test site shows matured bone formation, with rich vasculature. Abundant collagen fibre bundles can be seen. No multinucleated giant cells present. Minimal membrane remnants seen. Epithelial growth was also seen along with new bone formation in all the specimens. Compared to specimens at 3 months, fewer scaffold remnants were seen at the end of 6 months. *(Figure 3)*

**Figure 3.**
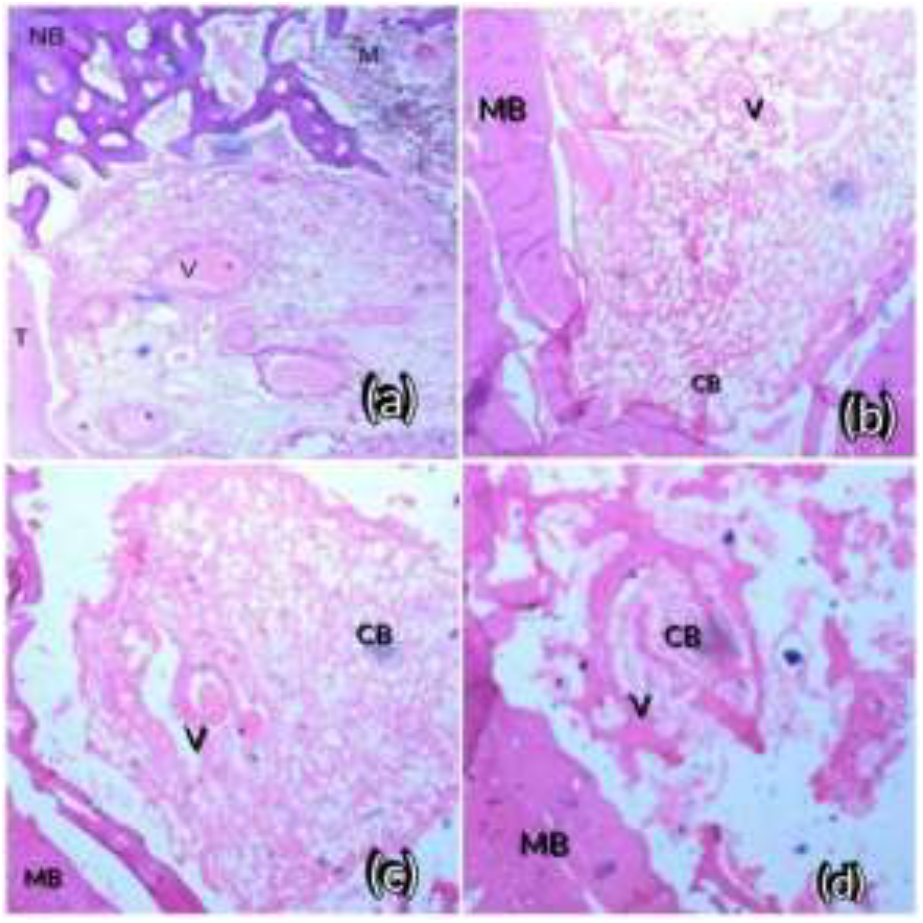
*T Tooth NB new bone V vasculature M membrane remnants* (a) 3 months histological specimen of test group; (b) 6 months histological specimen of test group; (c) 3 months histological specimen of control group; (d) 6 months histological specimen of control group

**Figure 4.**
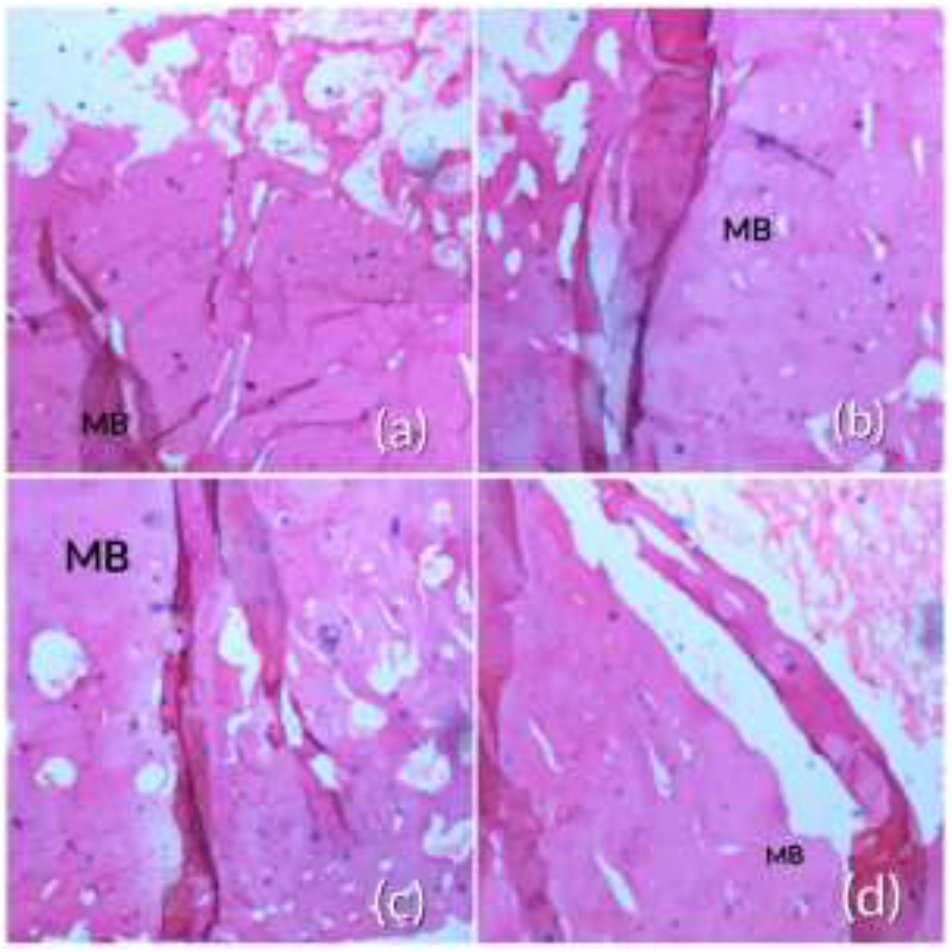
Analysing bone regeneration(a) Histological bone regeneration analysis of test sites at 3 months; (b) Histological bone regeneration analysis of test sites at 6 months; (c) Histological bone regeneration analysis of control sites at 3 months; (d) Histological bone regeneration analysis of control sites at 6 months

**Figure 5.**
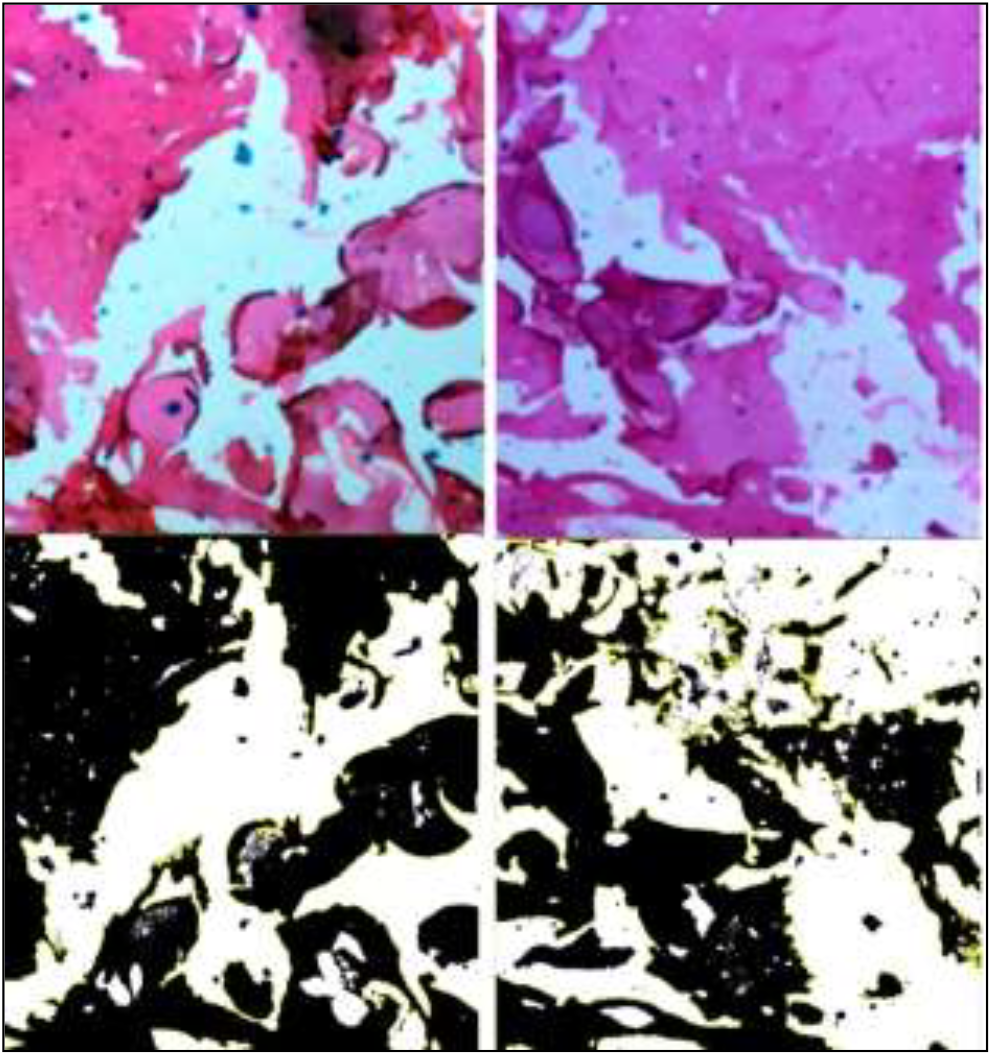
Mineralized tissue volume analysis

**Figure 6.**
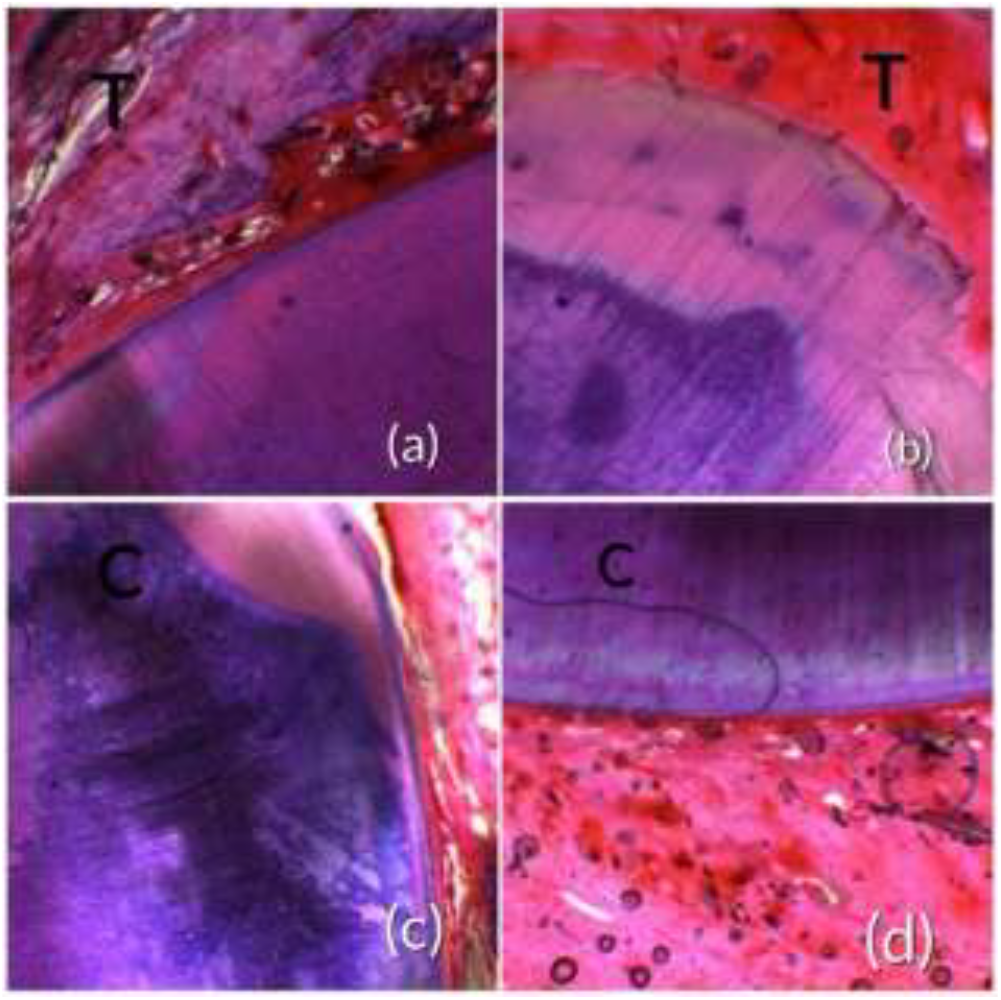
Analysing membrane remnants**(a)** test specimen at 3 months showing abundant membrane remnants (white areas) (b) test specimen at 6 months showing no membrane remnants; (c) 28 control specimen at 3 months showing abundant membrane remnants; (d)control specimen at 6 months showing less number of membrane remnants

The three and six months histological specimen of the control site showed similar findings to test group but new bone formation was lesser than in test group. *(Figure 3)*

### Mineralized Tissue Volume Analysis

The MTV values across T and C groups were 58.32±32.66 & 78.19±29.21 and 52.34±24.34 & 71.44±35.28 respectively at 3 and 6-months. There was a significant intergroup difference among these groups at 6-months. *(p=0*.*003)(Figure 5)*

### Analysing Membrane Remnants

Test specimen at 3 months showing abundant fibrovascular membranous remnants (white areas). Test specimen at 6 months showing no fibrovascular membranous remnants.

Control specimen at 3 months showing abundant fibrovascular membranous remnants. Control specimen at 6 months showing a smaller number of fibrovascular membranous remnants

The membrane degradation values across T and C groups were 60.19±23.70 & 66.69±14.20 and 97.54±16.67 & 95.25±16.56respectively at 3 and 6-months. Highly significant difference was seen for membrane degradation in test group *(p≤0*.*001**)* compared to control group. *(Table 1) (Figure 6)*

### Intergroup comparison of Recession depth (in mm) at various time intervals

When the results of recession coverage were compared within the test group, an increase in recession coverage was observed from 3 months (3.34±1.82) to 6 months (2.43±1.89) with rhFGF2 growth factor impregnated collagen membrane, whereas in control group also there was an increase in recession coverage from 3 months (3.58±1.68) to 6 months (2.98±1.50). When the results of recession coverage in rhFGF2 (test) group and regular collagen membrane (control) group were compared, at baseline no significant difference was seen between the test and control groups *(p=0*.*14)*. Significant differences were observed between the test and the control groups for the root coverage from baseline to 3 months. *(p=0*.*008*)* From baseline to 6-months post-operative intervals, the root coverage was highly significant in the test group when compared to the control group. *(p=0*.*001**) (Table 2) (Figure 2)*

**Table 2.** Intergroup comparison of Recession depth (in mm) at various time intervals.

| Time | Groups | Mean and standard deviation | t-value | p-value |
| --- | --- | --- | --- | --- |
| **Baseline** | *Test* | 5.89±1.47 | 2.908 | 0.14<br>(NS) |
|  | *Control* | 5.57±1.52 |  |  |
| **3-months** | *Test* | 3.34±1.82 | 6.52 | 0.008* |
|  | *Control* | 3.58±1.68 |  |  |
| **6-months** | *Test* | 2.43±1.89 | 7.83 | 0.001** |
|  | *Control* | 2.98±1.50 |  |  |

*Independent t-test, $p \leq 0.05$ \*(significant), $p \leq 0.001$ \*\* (highly significant), $p \geq 0.05$ NS (Not significant).*

## DISCUSSION

In the present study we investigated the efficacy of FGF 2 impregnated collagen membrane in healing of gingival recession in beagle dogs histologically. Improvement in recession depth was seen clinically, histologically bone regeneration, mineralised tissue volume, membrane remnants were evaluated, which showed significant results in the test group.

The new bone formation was seen in both the groups but in test group statistically significant difference was seen when compared to the control group. The results were in par with study conducted by Toshie Nagayasu-Tanaka^11^ et al in beagle dogs who opinioned that FGF-2 enhances new tissue formation at the early regeneration phase, leading to enhanced formation of new bone and FGF-2 increases the expression of osteoblastic differentiation markers, osterix, ALP in the regenerated tissue. Similarly, a rabbit study by Sang-Hoon Lee^12^ et al confirmed that polymer collagen membrane can be used as a carrier of FGF-2 when enhanced early stage of new bone formation is required.

FGF-2 accelerated the proliferation of fibroblasts derived from the PDL and bone marrow for new tissue formation. FGF-2 increases the production of BMP-2, vascular endothelial growth factor (VEGF) and transforming growth factor-β1 expression in human bone marrow stromal cells ^13^ and VEGF in human PDL stem/progenitor cell lines^14^. Thus, endogenous FGF-2 expression increases, administrated FGF-2 was considered to trigger the regenerative reaction including the proliferation and production of other growth factors.

Experiments using in-vitro maintained endothelial cells and PDL cells revealed that FGF-2 stimulated production of vascular endothelial growth factor (VEGF) in PDL cells and they synergistically induced angiogenesis.^15^ For regeneration, angiogenesis is essential to supply oxygen and nutrients, and is also required for successful bone induction during osteogenesis.^16^ FGF-2 increased alkaline phosphatase activity, osteocalcin production, and bone nodule formation following the enhancements of the proliferation of osteogenic progenitor cells from bone marrow.^17^ FGF-2 also showed to up-regulate the BMP-2 expression in human bone marrow stromal cells. These results suggest that FGF-2 may increase a specific cell population with high expression of BMP-2, or osteogenic progenitors in the regenerated tissue of the defect. This can be supported by histological results of present study where specimen of the surgical site (test) at three months showed woven bone (NB) with rich vasculature along with abundant collagen bundles. Epithelial growth was also seen along with new bone formation in all the specimens.

FGF signalling plays essential roles in osteogenesis and dentinogenesis, and among FGFs, FGF2 is widely expressed in the cells of odontoblast and osteoblast lineages and has been identified as a potent regulator of mineralization in-vivo and in-vitro.^18,19^ Yohei Nakayama et al., studied the efficacy of Fibroblast growth factor 2 mineralization-associated genes in osteoblast-like cell and concluded that, in human osteoblast-like Saos2 cells, FGF2 increased the expression of the osteopontin (SPP1, 16.7-fold), interleukin-8 (IL8, 6.4-fold) and IL11 (4.8-fold) genes. These results suggest that FGF2 might be a crucial regulator of mineralization and bone formation.^20^ Studies suggested that the effects of FGF signaling were depended on the stage of osteoblast maturation. In immature osteoblasts FGF signaling induced proliferation leading to increased osteogenesis in long term, whereas in mature osteoblasts FGF signaling inhibited differentiation and mineralization.^21, 22^ These evidences were consistent with the present study where a significant difference was seen in mineralized tissue volume in FGF 2 test group.

If the collagen membrane dissolves quickly, clinical treatment goals will not be achieved and regeneration will be unpredictable.^23^ An ideal barrier should resorb gradually over time.^24^ It has been suggested that these membranes must stay physically and mechanically intact for at least 4–6 weeks for regenerative therapy to be successful.^25^ The present study showed membrane remnants after 3 months suggesting its slow resorption rate which is helpful for bone regeneration. After 6 months, test specimen showed no fibrovascular membranous remnants, showing complete biodegradation or resorption of the scaffold. This was in accordance with a study conducted on differential biodegradation kinetics of collagen membranes for bone regeneration where native and defatted types I/III collagens were also almost completely resorbed after 12 weeks. ^26^

Periodontitis in dogs arises naturally from gingivitis with the aging process, but the time of onset is unpredictable and the defects are inconsistent. Therefore, in the present study the recession defect was created using bone chisels to expose the root surface. The main drawback of many periodontal procedures is the healing mechanism i.e. repair by formation of long junctional epithelium.^27, 28^ The test group showed statistically greater increase in root coverage at 3 months compared to the control group because FGF-2 shows early formation of connective tissue extending from the existing PDL seems to contribute to inhibiting down-growth of gingival epithelial tissue.^29, 30^ This was in acceptance with another preliminary human study where rhFGF-2-impregnated collagen membranes showed promising results in terms of increasing the wKG and recession coverage.^8^ Similarly, a study by Shujaa Addin et al., reported that rhFGF-2 in gelatin/β-TCP sponges exhibits an increased potential to support periodontal wound healing/regeneration in canine recession-type defects.^31^ In contrast to the present study, another animal study where FGF 2 in combination with β-TCP was used to evaluate the root coverage in dogs however the results showed enhanced formation of new bone and cementum without significant root coverage in this dog model.^32^ A sandwich membrane composed of a collagen sponge scaffold and gelatin microspheres containing basic fibroblast growth factor (bFGF) in a controlled-release system was developed by Taka Nakara et al which showed successful regeneration of the periodontal tissues in a short period of time.^33^

Human trials with larger sample size should be conducted to compare the equivalency in healing and regeneration. Other histological findings like true regeneration with cementum formation by FGF 2 should also be evaluated. Since the histological evidence showed bone formation, its efficacy in intra bony defects should also be assessed in the further studies.

## CONCLUSION

The three months and six months specimens of the test (rhFGF2 membrane) group has shown good amount of bone regeneration and soft tissue healing with limited membrane remnants at the site, when compared to the control group. The clinical recession depth was more in the test group(rhFGF2) than in the control group from baseline to three months and six months. These results prove that the rhFGF2 growth factor impregnated collagen membrane is highly efficient in treating gingival recession, thus improving the periodontal health.

## Notes

### Competing Interest Statement

The authors have declared no competing interest.

### Funding

- Biotechnology Ignition Grant (BIG), 16TH Cycle from Biotechnology Industry Research Assistance Council (BIRAC), INDIA: BIRAC/IKP01214/BIG-16/20

## REFERENCE

1. Pradeep K, Rajababu P, Satyanarayana D, Sagar V. Gingival recession: review and strategies in treatment of recession. Case Rep Dent. 2012;2012:563421. doi: 10.1155/2012/563421. Epub 2012 Oct 2. PMID: 23082256; PMCID: PMC3467775.

2. The etiology and prevalence of gingival recession Moawia M. Kassab, D.D.S. M.S.; Robert E. Cohen, D.D.S., Ph.D.

3. Madhuri SV. Membranes for periodontal regeneration. Int. J. Pharm. Sci. Invent. 2016 Oct;5(1):19–24.

4. Murakami, S. (2011). Periodontal tissue regeneration by signaling molecule (s): What role does basic fibroblast growth factor (FGF-2) have in periodontal therapy? Periodontology 2000, 56, 188–208. 10.1111/j.1600-0757.2010.00365.x

5. Outcomes of regenerative treatment with rhPDGF-BB and rhFGF-2 for periodontal intra-bony defects: a systematic review and meta-analysis

6. Urakami S, Takayama S, Kitamura M, Shimabukuro Y, Yanagi K, Ikezawa K, et al. Recombinant human basic fibroblast growth factor (bFGF) stimulates periodontal regeneration in class II furcation defects created in beagle dogs. J Periodontal Res 2003;38:97-103.n

7. Momose T, Miyaji H, Kato A, Ogawa K, Yoshida T, Nishida E, Murakami S, Kosen Y, Sugaya T, Kawanami M. Collagen Hydrogel Scaffold and Fibroblast Growth Factor-2 Accelerate Periodontal Healing of Class II Furcation Defects in Dog. Open Dent J. 2016 Jul 29;10:347–59. doi: 10.2174/1874210601610010347. PMID: 27583044; PMCID: PMC4974830.

8. Efficacy of recombinant human fibroblast growth factor 2 impregnated absorbable collagen membrane in the treatment of Miller’s Class I and II gingival recession defects Preliminary results from the first in human clinical trial

9. Struillou X, Boutigny H, Soueidan A, Layrolle P. Experimental animal models in periodontology: a review. The open dentistry journal. 2010;4:37.

10. Experimental Animal Models in Periodontology: A Review

11. Action Mechanism of Fibroblast Growth Factor-2 (FGF-2) in the Promotion of Periodontal Regeneration in Beagle Dogs

12. The role of rhFGF-2 soaked polymer membrane for enhancement of guided bone regeneration

13. Farhadi J, Jaquiery C, Barbero A, Jakob M, Schaeren S, Pierer G, et al. Differentiation-dependent upregulation of BMP-2, TGF-beta1, and VEGF expression by FGF-2 in human bone marrow stromal cells

14. Kono K, Maeda H, Fujii S, Tomokiyo A, Yamamoto N, Wada N, et al. Exposure to transforming growth factor-β1 after basic fibroblast growth factor promotes the fibroblastic differentiation of human periodontal ligament stem/progenitor cell lines. Ce

15. Yanagita M, Kojima Y, Kubota M, Mori K, Yamashita M, Yamada S, et al. Cooperative effects of FGF-2 and VEGF-A in periodontal ligament cells. J Dent Res. 2014; 93: 89–95. doi: 10.1177/0022034513511640 PMID: 24186558

16. ang TD, Salim A, Xia W, Nacamuli RP, Guccione S, Song HM, et al. Angiogenesis is required for successful bone induction during distraction osteogenesis. J Bone Miner Res. 2005; 20: 1114–1124. doi: 10.1359/JBMR.050301 PMID: 15940364

17. Pri-Chen S, Pitaru S, Lokiec F, Savion N. Basic fibroblast growth factor enhances the growth and expression of the osteogenic phenotype of dexamethasone-treated human bone marrow-derived bone-like cells in culture. Bone. 1998; 23: 111–117. doi: 10.1016/S8756-3282(98)00087-8 PMID: 9701469

18. Roberts-Clark DJ, Smith AJ. Angiogenic growth factors in human dentine matrix. Archives of oral biology. 2000; 45(11):1013–1016

19. Madan AK, Kramer B. Immunolocalization of fibroblast growth factor-2 (FGF-2) in the developing root and supporting structures of the murine tooth. Journal of molecular histology. 2005; 36(3): 171–178. [PubMed: 15900407]

20. Fibroblast growth factor 2 and forskolin induce mineralization-associated genes in two kinds of osteoblast-like cells Yohei Nakayama 1, 3

21. Miraoui H, Marie PJ. Fibroblast growth factor receptor signaling crosstalk in skeletogenesis. Science signaling. 2010; 3(146):re9. [PubMed: 21045207]

22. Marie PJ. Fibroblast growth factor signaling controlling bone formation: an update. Gene. 2012; 498(1):1–4. [PubMed: 22342254]

23. An, Y.-Z.; Kim, Y.-K.; Lim, S.-M.; Heo, Y.-K.; Kwon, M.-K.; Cha, J.-K.; Lee, J.-S.; Jung, U.-W.; Choi, S.-H. Physiochemical properties and resorption progress of porcine skin-derived collagen membranes: In vitro and in vivo analysis. Dent. Mater. J. 2018, 37, 332–340

24. Calciolari, E.; Ravanetti, F.; Strange, A.; Mardas, N.; Bozec, L.; Cacchioli, A.; Kostomitsopoulos, N.; Donos, N. Degradation pattern of a porcine collagen membrane in an in vivo model of guided bone regeneration. J. Periodontal Res. 2018, 53, 430–439.

25. Wong, C.; Yoganarasimha, S.; Carrico, C.; Madurantakam, P. Incorporation of Fibrin Matrix into Electrospun Membranes for Periodontal Wound Healing. Bioengineering 2019, 6, 57

26. Differential Biodegradation Kinetics of Collagen Membranes for Bone Regeneration Manuel Toledano

27. Trombelli L. Periodontal regeneration in gingival recession defects. P

28. Zucchelli G, Mele M, Mazzotti C, Marzadori M, Montebugnoli L, De Sanctis M. Coronally advanced flap with and without vertical releasing incisions for the treatment of multiple gingival recessions: a comparative controlled randomized clinical trial. Journal of Periodontology 2009; 80(7):1083–94.

29. Cha JK, Sun YK, Lee JS, Choi SH, Jung UW. Root coverage using porcine collagen matrix with fibroblast growth factor-2: a pilot study in dogs. Journal of Clinical Periodontology 2017; 44(1):96–103.

30. T Kao, S. Murakami, OR Beirne. The use of biologic mediators and tissue engineering in dentistry. Periodontology 2000 2009; 50(1):127–153.

31. Biodegradable gelatin/beta-tricalcium phosphate sponges incorporating recombinant human fibroblast growth factor-2 for treatment of recession-type defects: A split-mouth study in dogs

32. shii Y, Fujita T, Okubo N, Ota M, Yamada, S, Saito A. (2013) Effect of basic fibroblast growth factor (FGF-2) in combination with beta tricalcium phosphate on root coverage in dog. Acta Odontol Scand 2013;71:325–32

33. Nakahara T, Nakamura T, Kobayashi E, Inoue M, Shigeno K, Tabata Y, Eto K, Shimizu Y. Novel approach to regeneration of periodontal tissues based on in situ tissue engineering: effects of controlled release of basic fibroblast growth factor from a sandwich membrane. Tissue Eng. 2003 Feb;9(1):153–62. doi: 10.1089/107632703762687636. PMID: 12625964.

